# Sign epistasis can be absent in multi-peaked landscapes with neutral mutations

**DOI:** 10.1101/2025.09.30.679656

**Authors:** Dmitry N. Ivankov, Evgenii M. Zorin

**Author notes:** Author for correspondence: Dmitry N. Ivankov, Center for Molecular and Cellular Biology, Moscow, Russia.

## Abstract

Fitness landscapes provide a rigorous mathematical framework for analyzing evolutionary dynamics, including the study of epistasis, the main obstacle to predicting phenotype from genotype. In 2011, Poelwijk *et al*. formulated a foundational theorem stating that in any multi-peaked fitness landscape, “at least two mutations exhibit reciprocal sign epistasis” (Poelwijk *et al*., *J. Theor. Biol*., 272:141). The proof relied on the implicit assumption that neutral mutations are absent, commonly accepted in theoretical studies in evolutionary biology.

In this study, we extend Poelwijk *et al*.’s analysis by incorporating genotypes with equal fitness, specifically, accounting for neutral mutations. We demonstrate that when neutral mutations are considered, conventional pairwise reciprocal sign epistasis (RSE) may be entirely absent from a multi-peaked landscape. Instead, RSE is guaranteed only when considering “distant” RSE defined through composite mutations, wherein groups of mutations are treated collectively across all their possible combinations.

Applying these concepts to empirical fitness landscapes faces a practical limitation: phenotypic measurements contain experimental noise, making mutational effects statistically indistinguishable from zero. Under such conditions, statistically significant detection of RSE in multi-peaked landscapes may be impossible even when composite mutations are considered.

Theoretically, our findings imply that in the presence of neutral mutations, compensatory mutations in a multi-peaked fitness landscape need not be adjacent; rather, compensation can occur following one or more neutral steps along an evolutionary path. Practically, in real-world scenarios where fitness measurements contain uncertainty, there may be a fundamental technical limitation to detecting RSE in a statistically significant manner within multi-peaked landscapes.

## Introduction

Fitness landscapes (Wright, 1931) represent a convenient and straightforward mathematical concept for systematically studying various evolutionary phenomena, including epistasis, the dependence of a mutation’s effect on its genetic background (Fisher, 1918; Smith, 1970). Epistasis plays a crucial role in evolution (Kondrashov and Kondrashov, 2001; Weinreich *et al*., 2005; Breen *et al*., 2011; Weinreich *et al*., 2013; Weinstein *et al*., 2023) and is the primary obstacle to accurately predicting fitness or phenotype from genotype (Zhou *et al*., 2022). It is also critical in pathogenic deviations (Nagel, 2005; Ang *et al*., 2023) and contributes to diseases such as Alzheimer’s disease (Combarros *et al*., 2009; Lundberg *et al*., 2023) and autoimmune disorders (Rose and Bell, 2012). Therefore, understanding the theoretical and empirical rules linking epistasis to other evolutionary phenomena is of great importance.

One key type of epistasis is reciprocal sign epistasis (RSE), which occurs when two mutations exhibit compensatory effects in a specific genetic context (Poelwijk *et al*., 2011). Specifically, each mutation partially, fully, or excessively counteracts the effect (deleterious or beneficial) of the other (Poelwijk *et al*., 2011; Kvitek and Sherlock, 2011). RSE drives compensatory evolution (Mammano *et al*., 2000; Jucovic and Hartley, 1996; Baresić *et al*., 2010; Ivankov *et al*., 2014) and is associated with traversing fitness valleys, making it relatively rare in proteins (Baresić *et al*., 2010). RNA molecules provide the clearest examples of RSE: Watson-Crick pairings evolve between A:U and G:C via intermediates G:U (more common) or A:C (rarer) (Meer *et al*., 2010).

In 2011, Poelwijk *et al*. formulated a fundamental theorem stating that “Reciprocal sign epistasis is a necessary condition for multi-peaked fitness landscapes” (Poelwijk *et al*., 2011). For brevity, we refer to this as “Poelwijk’s theorem” throughout this paper. The theorem guarantees that along the shortest pathways between any two peaks in a multi-peaked landscape, “at least two mutations exhibit reciprocal sign epistasis” (Poelwijk *et al*., 2011). The proof was based on a common and widely accepted assumption in theoretical evolutionary biology that all genotypes in the landscape have distinct fitness values.

In this study, we extend the analysis of Poelwijk *et al*. (2011) by considering the presence of equal-fitness genotypes, specifically neutral mutations (Kimura, 1968; Ohta, 1973) in multi-peaked fitness landscapes. We demonstrate that in landscapes containing neutral mutations, only general RSE – uniting conventional (pairwise) RSE and distant RSE – can be guaranteed. Distant RSE refers to cases where composite mutations, i.e., groups of single mutations analyzed collectively in all possible combinations, are considered (Zorin *et al*., 2022). When switching focus to the practical task of detecting RSE in empirical landscapes, we encounter the challenge that phenotypic measurements inherently include some degree of uncertainty. Under these conditions, even considering composite mutations does not ensure statistically significant detection of general RSE.

Our findings carry both theoretical and practical implications. First, in multi-peaked landscapes containing neutral mutations, compensatory mutations need not be adjacent along evolutionary paths. Second, in real-world scenarios where fitness measurements are inevitably subject to uncertainty, statistically significant detection of general reciprocal sign epistasis (or sign epistasis) – whether analyzing single or composite mutations – may be technically impossible within multi-peaked landscapes.

## Results

### Reciprocal sign epistasis and Poelwijk’s theorem

In 2011, Poelwijk *et al*. introduced the highly insightful term “reciprocal sign epistasis” (RSE) to describe compensatory evolutionary events at the level of single mutations (see Definition 4 in (Poelwijk *et al*., 2011)). This is a special case of sign epistasis (see Definition 3 in (Poelwijk *et al*., 2011)). Poelwijk *et al*. (2011) proved a foundational theorem that “reciprocal sign epistasis is a necessary condition for multi-peak landscapes”.

The proof relies on the implicitly stated standard assumption in theoretical evolutionary biology that all genotypes in the fitness landscape have distinct fitness values. For the reader’s convenience, a verbatim reproduction of Poelwijk’s theorem and its proof is provided in the Supplementary Materials. An illustration of the theorem’s application is shown in Fig. 1.

**Fig. 1.**
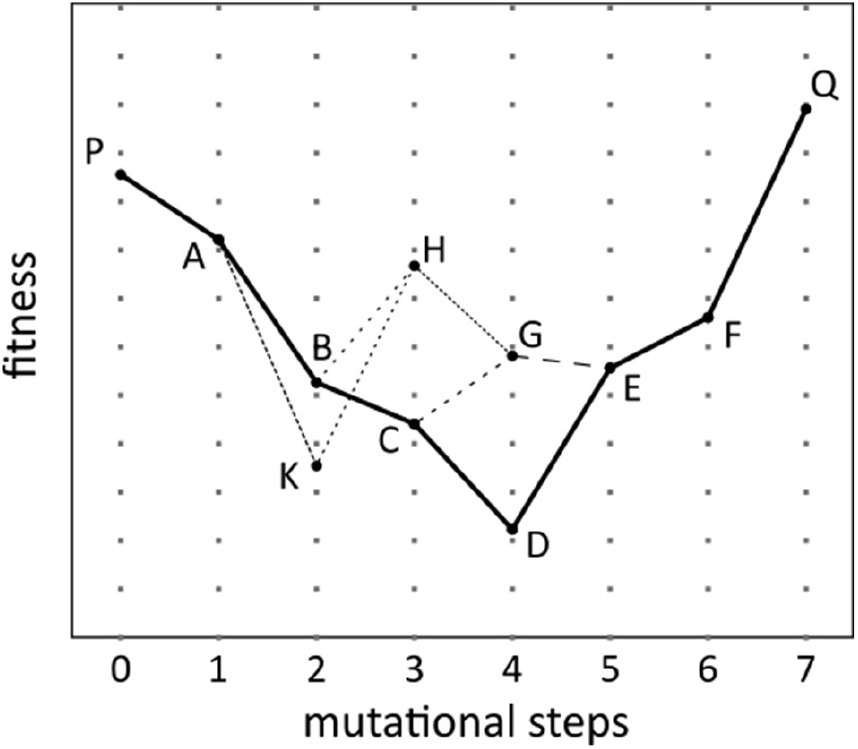
Illustration of the proof of the Poelwijk’s theorem with numbers in parentheses referencing the steps from the proof. Adapted from Poelwijk *et al*. (2011). – The existence of the two maxima, P and Q, is established (step (1)). – A random direct path PABCDEFQ of length 7 from P to Q is considered (step (2)). – This path contains one minimum D between genotypes C and E (step (3)). – The order of mutations C→D and D→E around the minimum D is reversed, becoming C→G and G→E (mutations C→G and D→E are the same mutation applied in different genetic contexts C and D; mutations G→E and C→D are the same mutation applied in different genetic contexts G and C), and the new path PABCGEFQ is considered (step (4)). – The minimum in the new path is found between new genotypes B and G, at point C (steps (5) and (5b)), so we proceed to step (4). – The order of mutations B→C and C→G around the new minimum C is reversed, becoming B→H and H→G, and the new path PABHGEFQ is considered (step (4)). – The minimum in the new path is found between new genotypes A and H, at point B (steps (5) and (5b)), so we proceed to step (4). – The order of mutations A→B and B→H around the new minimum B is reversed, becoming A→K and K→H, and the new path PAKHGEFQ is considered (step (4)). – The minimum in the new path, is found between the same genotypes A and H as previous minimum B, at point K, so the RSE is detected in the group of genotypes A, B, H, and K (steps (5) and (5a)).

### Scenarios of fitness equality

The subsequent analysis addresses two key questions: (1) the existence of RSE in multi-peaked landscapes containing genotypes with equal fitness values, and (2) how measurement errors can affect the detection of RSE in experimental fitness landscapes.

The occurrence of equal-fitness genotypes in fitness landscapes can be categorized into two distinct scenarios: (1) when equal-fitness genotypes are distant from each other in the landscape, and (2) when they are adjacent (i.e., connected by neutral mutations).

We analyze these cases separately, beginning with distant equal-fitness genotypes. We then examine landscapes with neutral mutations and discuss the practical implications of our results for experimental settings, where mutational effects may be statistically indistinguishable from zero.

### Reciprocal sign epistasis is a necessary condition for multi-peaked landscapes having distant equal-fitness genotypes

When two distant genotypes along the pathway between two peaks have equal fitness and serve as minima (Fig. 2), RSE remains a necessary condition in multi-peaked landscapes. The extension of Poelwijk’s theorem for this case is as follows:

**Fig. 2.**
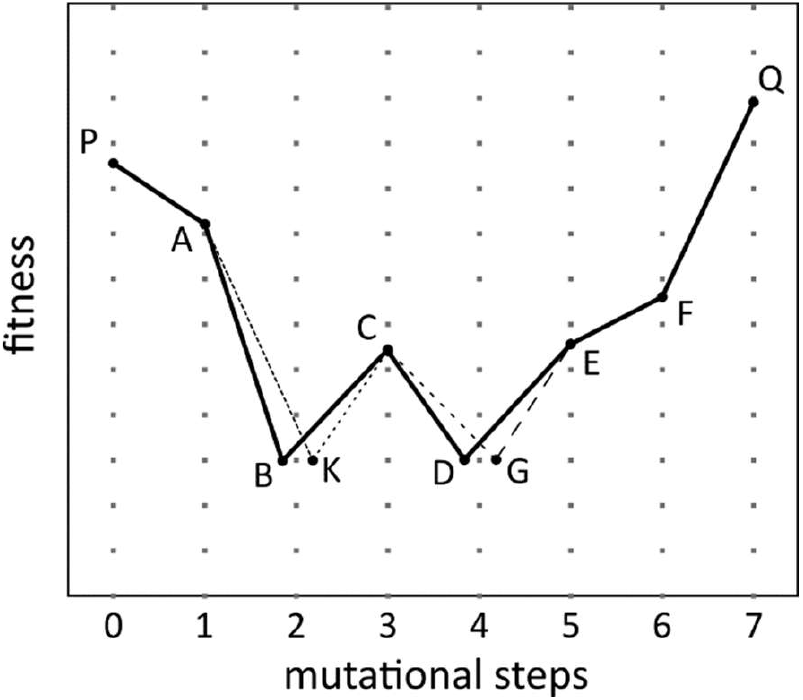
Example of a multi-peaked landscape with distant genotypes having equal fitness. Genotypes B, K, D, and G share the same fitness. Genotypes B and K are two mutational steps from genotype P, while genotypes D and G are four mutational steps away from P. To avoid overlapping genotypes in the picture, B, K, D, and G are slightly shifted from their exact vertical positions.

#### Theorem 1.

In a bi-allelic *L*-loci system, reciprocal sign epistasis is a necessary condition for the existence of two peaks at a distance *L* in a fitness landscape that can contain distant equal-fitness genotypes.

The original proof (Poelwijk *et al*., 2011) requires only a minor modification to accommodate this scenario: if multiple equal-fitness minima exist along the randomly selected pathway at step (3) randomly select one minimum. The modified proof, along with the general formulation of Theorem 1, is provided in the Supplementary Materials.

### Reciprocal sign epistasis and sign epistasis can be absent in multi-peaked landscapes with neutral mutations

When genotypes with equal fitness are neighbors in sequence space, neutral mutations are present. Consider the minimal multi-peaked fitness landscape that shows such a landscape may not contain conventional (i.e., pairwise) RSE. This landscape represents a combinatorially complete dataset for three loci, each with two alleles, where local maxima occur at the wild-type genotype P and the triple mutant Q. All intermediate single and double mutants – A, B, C, D, E, and F – have equal and low fitness (Figure 3). Note that this differs from simply adding a universally neutral third site to the conventional RSE scenario (see right panel of Fig. 1 in (Poelwijk et al., 2011)) because the neutrality of each mutation depends on the genetic context. As can be seen by manual inspection of all six two-dimensional combinatorially complete datasets (PACB, PAED, PCEF, ABDQ, CBFQ, and EDFQ), there are no instances of either RSE or sign epistasis in this landscape.

**Fig. 3.**
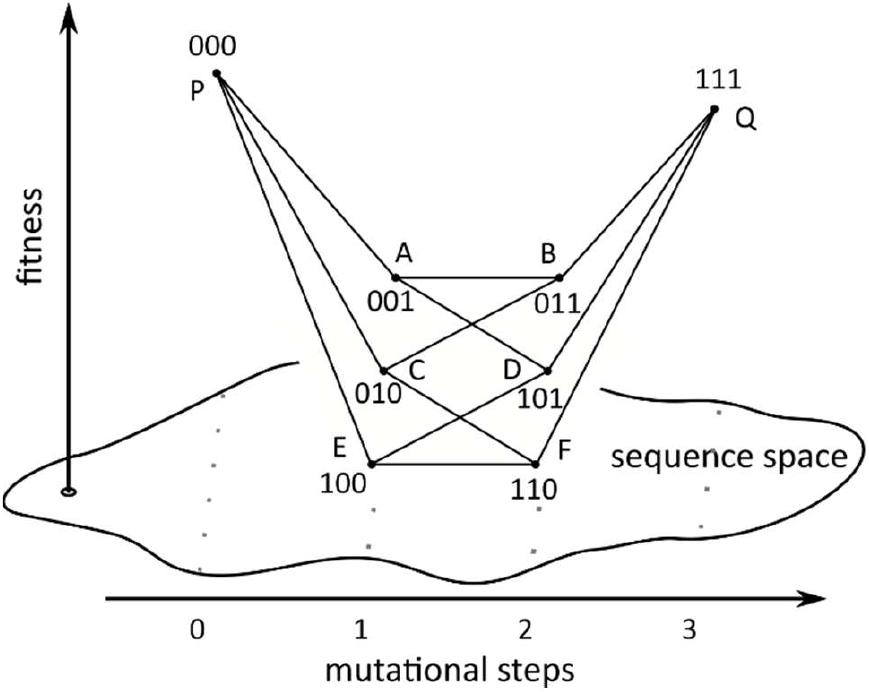
A landscape showing a combinatorially complete dataset for three loci, each with two alleles. The fitness values of the single and double mutants, that is, of genotypes A, B, C, D, E, and F, are equal to each other.

Sign epistasis occurs when the sign of a mutation’s fitness effect depends on the genetic background (Definition 3 in (Poelwijk *et al*., 2011)). If the effect of a mutation in one of the two backgrounds is zero, this might be considered as a “weak sign epistasis”: the mutation has a positive (or negative) effect in one background and a non-positive (or non-negative) effect in the other. A “weak RSE” version of RSE could be formulated analogously. Under these relaxed formulation, Poelwijk’s theorem would remain valid even in landscapes containing neutral mutations. However, we adhere to the strict definitions of both sign and reciprocal sign epistasis given by Poelwijk *et al*. (2011) (Definition 3 and 4).

Thus, Poelwijk’s theorem cannot be extended to the general case of multi-peaked landscapes containing neutral mutations. The original theorem remains valid only in the absence of neutral mutations.

### Only general RSE is guaranteed for multi-peaked landscapes having neutral mutations

Consider two fitness peaks, P and Q, separated by lower-fitness genotypes (Figure 3). Along pathways between them, fitness initially decreases in all directions before being compensated upon reaching genotype Q. In the landscape shown in Figure 3, this compensation is delayed. For example, on the path PEFQ, the first mutation occurs at the first locus (P: 000 → E: 100), followed by a neutral second mutation at the second locus (E: 100 → F: 110). The final mutation at the third locus compensates for the initial fitness decrease, but only in the context of genotype F (110). Importantly, this does not constitute immediate compensation for the first mutation, as the step from E (100) to D (101) is neutral (note that no instances of immediate compensation occur anywhere in this landscape). Thus, compensation occurs through a “composite” mutation (E: 100 → Q: 111) involving both the second and third loci together (Zorin *et al*., 2022). RSE in this landscape is observed only distantly, i.e., when considering combinations of mutations rather than individual ones, leading to the following theorem:

#### Theorem 2.

In a bi-allelic *L*-loci system, general RSE (uniting conventional RSE and distant RSE) is a necessary condition for the existence of two peaks separated by distance *L* in the fitness landscape containing neutral mutations.

Proof strategy for Theorem 2 is to proceed iteratively by dimensionality reduction (illustrated using a four-loci combinatorially complete dataset):

1. Identify two peaks, e.g., genotypes “0000” (peak P) and “1111” (peak Q).
2. Select the lower peak, say “0000” (P). All its neighbors (“0001”, “0010”, “0100”, “1000”) have lower fitness.
3. Choose in the considered landscape any neighbor of P, e.g., “0001”.
4. Find the genotype most distant from “0001” in the considered landscape, which is “1110”.
5. Case 1: If fitness(“1110”) < fitness(“0000”), then distant RSE exists among “0000”, “0001”, “1110”, and “1111”.
6. Case 2: Otherwise, reduce to a three-loci landscape (from “0000” to “1110” with all shortest pathways between them) and repeat the procedure from step 3.

The process terminates when either distant RSE is found or we reach genotype two steps away from peak P (e.g., W: “1100”). At this point, conventional (pairwise) RSE is guaranteed, as all direct neighbors of P – including those “0100” and “1000” between W and P – have lower fitness. A more formal proof of Theorem 2 is provided in the Supplementary Materials.

### In experimentally measured landscapes both conventional RSE and distant RSE may not be detected

We have just examined theoretical extensions of Poelwijk’s theorem that account for both distant genotypes with equal fitness values and neutral mutations. The next step is to consider how the proofs of Poelwijk’s theorem and its extensions presented here can be applied to practically detect RSE in experimental fitness landscapes. This is straightforward because all proofs are formulated “instrumentally” as search algorithms.

In theory, we assume phenotypes can be measured with infinite precision, so they are either distinct or exactly equal. Consequently, if two genotypes with equal phenotypes are adjacent in the landscape, the mutation between them is either beneficial, deleterious, or neutral. In experimental landscapes, however, we encounter reality where fitness (and its proxies) is estimated from measurements subject to error. Thus, each fitness value should be reported with its associated standard deviation.

If all genotypes in a multi-peaked empirical landscape have statistically distinct fitness values, then applying Poelwijk’s theorem proof as a computer algorithm will perfectly detect RSE.

If a multi-peaked landscape contains distant genotypes with statistically indistinguishable fitness values but no neutral mutations, then applying Theorem 1’s proof will also perfectly detect RSE. This is because the presence of only distant equal-fitness genotypes (without neutral mutations) ensure that adjacent genotypes along any path have distinct fitness values, making Theorem 1 and its proof directly applicable.

For mutations that are statistically indistinguishable from neutral, consider the measurement of a hypothetical landscape shown in Figure 4, representing a bi-allelic three-loci combinatorially complete dataset. Suppose the fitness values were measured with uncertainties such that, at a chosen *a priori* threshold of statistical significance, the single mutants A, C, and E are statistically indistinguishable from the double mutants B, D, and F, which in turn are statistically indistinguishable from the fitness of the triple mutant Q.

**Fig. 4.**
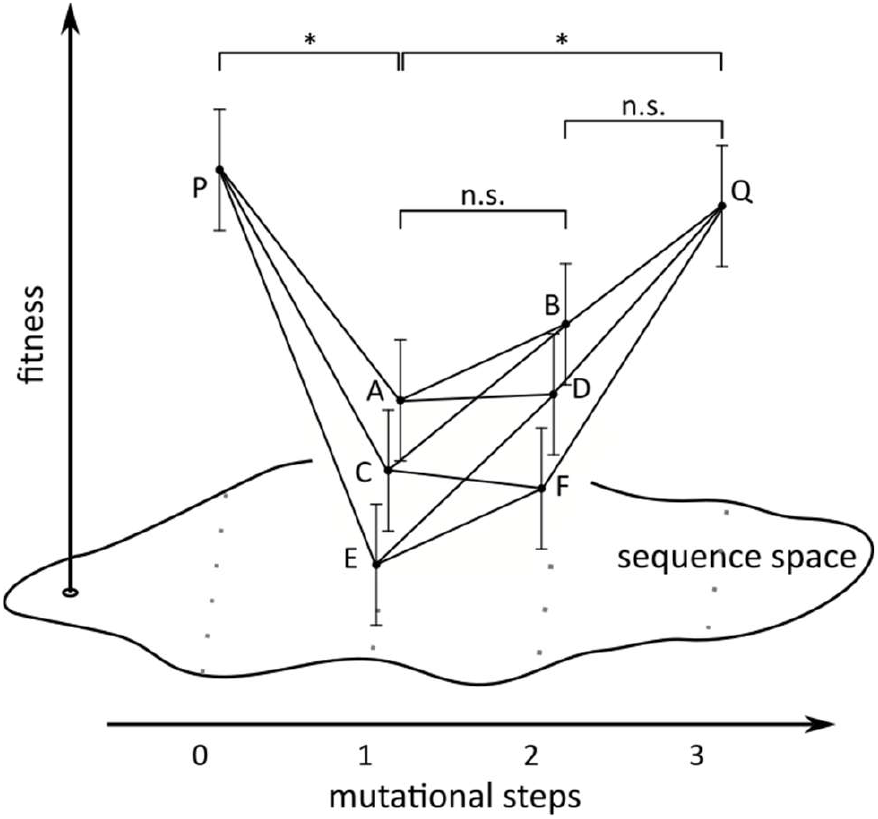
Example of a bi-allelic three-loci landscape where fitness values are measured with some error, indicated by vertical bars. Single mutants A, C, and E have identical fitness that is significantly lower (denoted by ‘*’) than the fitness of genotypes P and Q, but statistically indistinguishable (denoted by ‘n.s.’) from the fitness of double mutants B, D, and F, which in turn are statistically indistinguishable from the fitness of genotype Q.

First, note that at the chosen threshold of statistical significance, this landscape has more than one peak. Indeed, in Figure 4, genotype P is clearly a peak, as all its neighbors (A, C, E) have significantly lower fitness. A second peak also exists, as genotype Q has significantly higher fitness than genotypes A, C, and E, which form a valley between P and Q. However, since Q is statistically indistinguishable from B, D, and F, the exact location of this second peak is uncertain but must lie somewhere within the group {Q, B, D, F}.

(If the true fitness values were known, Q and B might share the highest fitness within this group, forming a quasi-peak; alternatively, B, D, and F might exceed Q, creating three separate peaks and making the landscape four-peaked overall. Such analysis, however, is beyond the scope of this paper. For our purposes, it is important that more than one peak is present in the entire landscape.)

Returning to RSE detection in Figure 4: manual inspection of all six two-dimensional combinatorially complete datasets (PACB, PAED, PCEF, ABDQ, CBFQ, EDFQ) reveals no statistically significant RSE or sign epistasis since mutations between single (A, C, E) and double (B, D, F) mutants are statistically indistinguishable from neutral. Even using the concept of composite mutations and examining the three remaining distant two-dimensional datasets (PAFQ, PCDQ, and PEBQ), no statistically significant distant RSE or sign epistasis is observed, since mutations from double mutants (B, D, F) to triple mutant Q are also statistically indistinguishable from neutral. This simple example illustrates the principal challenges in applying Poelwijk’s theorem and its proposed extensions to detect RSE and sign epistasis in experimentally measured fitness landscapes.

## Discussion

The relationship between types of epistasis and fitness landscape properties is a key question in evolutionary biology. The presence of RSE in multi-peaked landscapes where all genotypes have distinct fitness values – proven by Poelwijk *et al*. (2011) – has been the most notable example.

Here we extend Poelwijk’s analysis to show that RSE still occurs when distant genotypes with equal fitness values are present in the landscape. In landscapes with neutral mutations (Kimura, 1968; Ohta, 1973), however, only general RSE (combining conventional RSE and distant RSE) can be guaranteed, requiring the composite mutation approach (Zorin *et al*., 2022).

In experimental fitness landscapes, measurement error may render detection of any form of RSE or sign epistasis practically impossible.

Poelwijk *et al*.’s proof assumes distinct phenotypes for different genotypes, which holds theoretically. Practically, this requires sufficient experimental replicates to detect statistically significant differences between initially similar or even identical phenotypes.

The discreteness of experimental fitness landscapes is often artificial, arising from technical limitations that bin protein variants into discrete categories (Adkar *et al*., 2012). However, some traits are naturally discrete; for example, fitness measured as offspring number inherently produces discrete values. These are estimates of fitness; and, ideally, replicate measurements can determine both their average across experiments and associated standard deviation. However, such replicates are sometimes difficult or impossible to obtain: for instance, measurements of human fitness as offspring number cannot be replicated, at least for ethical reasons.

In practice, many phenotypes in experimental landscapes are measured only a few times, sometimes once, underscoring that research typically operates with imperfect phenotypic accuracy. Thus, the proofs of Poelwijk’s theorem and its proposed extensions represent distinct challenges from their practical application for detecting RSE in experimental fitness landscapes.

The framework of considering combinations of single mutations as a single “composite” mutations that was useful here, is described by Zorin *et al*. (2022). While it offers limited evolutionary insight, since the probability of multiple mutations at specific loci within one generation is prohibitively low (Smith, 1970), both single and multiple mutations represent just changes in protein chemical structure. Composite mutations enable analysis of adaptive landscapes at any distance, revealing distant epistasis alongside conventional epistasis and maximizing epistasis detection (Zorin *et al*., 2022). Leveraging efficient algorithm (Esteban *et al*., 2019) for identifying combinatorially complete datasets in genotype-phenotype maps generated by (quasi-)random mutagenesis (Sarkisyan et al., 2016; Pokusaeva et al., 2019), this approach dramatically expands the repertoire of such datasets and enables distant epistasis detection (Zorin *et al*., 2022).

Poelwijk *et al*. (2011) clearly did not intend composite mutations in their study, as they explicitly refer to “single mutations” throughout their paper. Moreover, we did not find a way to substitute composite mutations into the original proof. For example, it would be implausible to interpret “changing the order of mutations leading to the current minimum” (Poelwijk *et al*., 2011) as involving composite mutations.

It is also important to address analysis at the gene level, where each locus represents an entire gene rather than a single protein sequence position. Here, a “single mutation” denotes a transition between gene variants, potentially involving numerous amino acid substitutions, insertions, or deletions. However, such a mutation is not considered composite, as compositeness is defined by the analytical approach – specifically, probing all combinations of single mutations to identify distant combinatorially complete datasets in protein sequence space.

Additionally, note the symmetry among single and double mutants in Figure 3: all single mutants and double mutants share identical fitness values. This symmetry ensures that absence of reciprocal sign epistasis implies absence of sign epistasis, since non-reciprocal sign epistasis would be asymmetric.

To summarize, Poelwijk’s theorem establishes a key link between RSE and multi-peaked fitness landscapes in the theoretical framework where equal-fitness genotypes – and specifically neutral mutations – are absent. We extend Poelwijk’s theorem to landscapes with distant equal-fitness genotypes (Theorem 1). In the presence of neutral mutations, however, only general RSE (uniting conventional RSE and distant RSE) can be guaranteed (Theorem 2). Practically, detecting any type of RSE or sign epistasis may be impossible in landscapes with mutations statistically indistinguishable from neutral due to measurement limitations.

## Acknowledgements

We thank Dmitry Biba and the anonymous reviewer for their insightful and constructive comments and valuable recommendations.

