## Supplementary Materials for "Sign epistasis can be absent in multi-peaked landscapes with neutral mutations"

#### Reciprocal sign epistasis and Poelwijk's theorem

For the reader's convenience, we provide here a verbatim copy of Poelwijk's theorem and its proof (Poelwijk et al., 2011). The Theorem relies on the implicitly stated standard assumption in theoretical evolutionary biology that all genotypes in the fitness landscape have distinct fitness values.

**Poelwijk's theorem.** "In a bi-allelic *L* locus system, reciprocal sign epistasis is a necessary condition for the existence of two peaks at a distance *L* in the fitness landscape." (Poelwijk et al., 2011)

**Proof.**

"(1) There are two maxima at distance *L* in a bi-allelic *L* locus system.

(2) We consider a random direct path (length *L*) from one maximum to the other one.

(3) This path necessarily contains a minimum.

(4) We change the path, by reversing the order of the two mutations leading to and away from the minimum.

(5) Two cases are possible:

- (a) If the fitness minimum of the path is still located between the same mutations, then these mutations exhibit reciprocal sign epistasis.
- (b) If the fitness minimum is now located at another step along the path, go back to point (4), now for the two mutations around this new minimum.

(6) The repeating of steps (4) and (5) necessarily leads to breaking the loop via case (5a), because the maximization of the minimum along the path is necessarily bounded (by the value of the lowest peak)." (Poelwijk et al., 2011).

**End of proof.**

Poelwijk *et al.* provided three arguments (see them numbered in Appendix, p.144 of (Poelwijk *et al.*, 2011)) showing that their statement about two peaks in a bi-allelic *L*-loci system is equivalent to an analogous statement about multiple peaks in an *N*-allelic *L*-loci system. Therefore, the general formulation of Poelwijk's theorem is as follows:

**Poelwijk's theorem (general formulation).** "In a *N*-allelic *L* locus system, reciprocal sign epistasis is a necessary condition for the existence of multiple peaks in the fitness landscape." (Poelwijk et al., 2011)

### Reciprocal sign epistasis is a necessary condition for multi-peaked landscapes having distant equal-fitness genotypes

Here we provide formulation of Theorem 1, its proof (which is a slightly modified proof of Poelwijk's theorem), as well as the general formulation of Theorem 1.

**Theorem 1.** In a bi-allelic *L*-loci system, reciprocal sign epistasis is a necessary condition for the existence of two peaks at a distance *L* in a fitness landscape that can contain distant equal-fitness genotypes.

**Proof.**

(1) Assume there are two maxima at distance *L* in a bi-allelic *L*-loci system.

(2) Consider a random direct path (necessarily of length *L*) between the two maxima.

(3) This path necessarily contains at least one minimum. If multiple minima exist, choose one at random.

(4) Modify the path by reversing the order of the two mutations leading to and away from this minimum.

(5) Two cases arise:

(a) If the genotype replacing the current minimum has lower fitness than its neighboring genotypes on the path, reciprocal sign epistasis is detected. Note that neighboring genotypes always have different fitness values as neutral mutations are absent.

(b) Otherwise, find the new minimum along the path. If multiple minima exist, randomly choose one. Then return to step (4) for the mutations surrounding this new minimum.

(6) Repeating steps (4) and (5) must eventually break the loop via case (5a), since the minimum fitness along the path is bounded above by the value of the lowest peak.

**End of proof.**

Taking into account the three numbered arguments given in Appendix of (Poelwijk *et al.*, 2011), we can provide general formulation of Theorem 1:

**Theorem 1 (general formulation).** In an *N*-allelic *L*-loci system, reciprocal sign epistasis is a necessary condition for the existence of multiple peaks at a distance *L* in a fitness landscape that can contain distant equal-fitness genotypes.

### Only general RSE is guaranteed for multi-peaked landscapes having neutral mutations

Here we provide formulation of Theorem 2, its proof, as well as the general formulation of Theorem 2.

**Theorem 2.** In a bi-allelic *L*-loci system, general RSE (uniting conventional RSE and distant RSE) is a necessary condition for the existence of two peaks separated by distance *L* in the fitness landscape containing neutral mutations.

**Proof.**

(1) Assume two maxima at distance *L* in a bi-allelic *L*-loci system. Label the lower maximum as P and the higher as Q. If the two maxima have equal fitness, arbitrarily label them P and Q. Thus, $f(P) \leq f(Q)$. By redefining Q as M, we have:

$$f(P) \leq f(M).$$

(2) Select in the analyzed landscape any single mutant A of genotype P. Since P is a peak, then:

$$f(A) < f(P). \qquad [1]$$

(3) Identify genotype B maximally distant from genotype A in the analyzed landscape.

(4) Two cases arise:

(a) Case 1: If

$$f(B) < f(P), \qquad [2]$$

then general RSE occurs, because:

$$f(A) < f(P) \leq f(M) \text{ implies:}$$

$$f(A) < f(M); \qquad [3]$$

$$f(B) < f(P) \leq f(M) \text{ implies:}$$

$$f(B) < f(M). \quad [4]$$

Conditions [1], [2], [3], and [4] mean that genotypes P, A, B, and M form general RSE.

(b) Case 2: If $f(B) \geq f(P)$, relabel B as M, reset B, and analyze one-locus less twice smaller landscape containing only shortest paths between P and M by repeating procedure from step (2).

(5) Repeating steps (3) and (4) must terminate in case (4a) because each iteration reduces the landscape by one locus. Eventually, either (i) distant RSE is found in step (4a) or (ii) the system reduces to two loci, where the genotype M is just two steps away of peak P. Here conventional RSE is guaranteed in step (4a) since all neighbors of P (including two mutants between P and M) have fitness lower than P (which in turn lower than M).

**End of proof.**

Taking into account the three numbered arguments given in Appendix of (Poelwijk *et al.*, 2011), we can provide general formulation of Theorem 2:

**Theorem 2 (general formulation).** In an *N*-allelic *L*-loci system, general RSE (uniting conventional RSE and distant RSE) is a necessary condition for the existence of multiple peaks separated by distance *L* in the fitness landscape containing neutral mutations.
